# Investigating the combined association of BMI and alcohol consumption on liver disease and biomarkers: a Mendelian randomization study of over 90 000 adults from the Copenhagen General Population Study

**DOI:** 10.1101/303313

**Authors:** Alice R Carter, Maria-Carolina Borges, Marianne Benn, Anne Tybjærg-Hansen, George Davey Smith, Børge G Nordestgaard, Debbie A Lawlor

## Abstract

**Background:** Body mass index (BMI) and alcohol consumption are suggested to independently and interactively increase the risk of liver disease. We assessed this combined effect using factorial Mendelian randomization (MR).

**Methods:** We used multivariable adjusted regression and MR to estimate individual and joint associations of BMI and alcohol consumption and liver disease biomarkers (alanine aminotransferase (ALT) y-glutamyltransferase (GGT)) and incident liver disease. We undertook a factorial MR study splitting participants by median of measured BMI or BMI allele score then by median of reported alcohol consumption or *ADH1B* genotype (AA/AG and GG), giving four groups; low BMI/low alcohol (-BMI/-alc), low BMI/high alcohol (-BMI/+alc), high BMI/low alcohol (+BMI/-alc) and high BMI/high alcohol (+BMI/+alc).

**Results:** Individual positive associations of BMI and alcohol with ALT, GGT and incident liver disease were found. In the factorial MR analyses, considering the +BMI/+alc group as the reference, mean circulating ALT and GGT levels were lowest in the -BMI/-alc group (-2.32% (95% CI: −4.29, −0.35) and −3.56% (95% CI: −5.88; −1.24) for ALT and GGT respectively). Individuals with -BMI/+alc and +BMI/-alc had lower mean circulating ALT and GGT compared to the reference group (+BMI/+alc). For incident liver disease multivariable factorial analyses followed a similar pattern to those seen for the biomarkers, but little evidence of differences between MR factorial categories for odds of liver disease.

**Conclusions:** Consistent results from multivariable regression and MR analysis, provides compelling evidence for the individual adverse effects of BMI and alcohol consumption on liver disease. Intervening on both BMI and alcohol may improve the profiles of circulating liver biomarkers. However, this may not reduce clinical liver disease risk.

### Key Messages

- Individually, BMI and alcohol consumption causally increase circulating levels of biomarkers of liver injury
- Factorial MR provides a valid approach for investigating the joint effects of risk factors, less prone to bias from confounding and reverse causality compared with observational study designs.
- Results from multivariable and Mendelian randomization analyses suggest BMI and alcohol are most likely to act independently, rather than interactively on increasing levels of circulating liver function biomarkers

## Introduction

Risk of liver disease in the population has increased, and the age of first diagnosis has decreased in many high income countries in recent decades (1). A key contributor is likely to be the global obesity epidemic and the relationship between greater adiposity and nonalcoholic fatty liver disease (NAFLD) (2). Increasing, or sustained, unhealthy levels of alcohol consumption in some countries is also likely to have contributed (3).

Recent studies suggest that overweight and alcohol consumption positively interact (on a multiplicative scale) to increase the risk of liver pathology, suggesting the relative risk of liver disease in people who drink excessively and are overweight or obese is greater than would be expected if the two risk factors were independent (4). Consistent with these findings, several studies have found that greater body mass index (BMI) and greater alcohol consumption positively interact in relation to biomarkers of liver injury, such as alanine aminotransferase (ALT), aspartate aminotransferase (AST) and y-glutamyltransferase (GGT) (5, 6). By contrast, in a study of British women (~ 1.2 million women with 1811 occurrences of a first hospital admission or death from liver cirrhosis) there was no evidence of an interaction between BMI and alcohol in relation to cirrhosis (7).

Understanding the combined effects of BMI and alcohol on liver disease is important for developing preventive public health interventions and establishing the likely future burden of liver disease in populations with differing levels of high BMI and high alcohol consumption. Previous findings from observational studies might be explained by residual confounding, misclassification bias, or reverse causality. Mendelian randomization (MR), the use of genetic variants as instrumental variables for assessing the effect of modifiable exposures, is increasingly used to establish causal relationships between risk factors and outcomes. MR can be used in a factorial design when considering how two or more risk factors may work together to influence an outcome, comparable to a factorial randomised control trial (RCT) design (**Figure 1** adapted from; (8)).

**Figure 1A.**
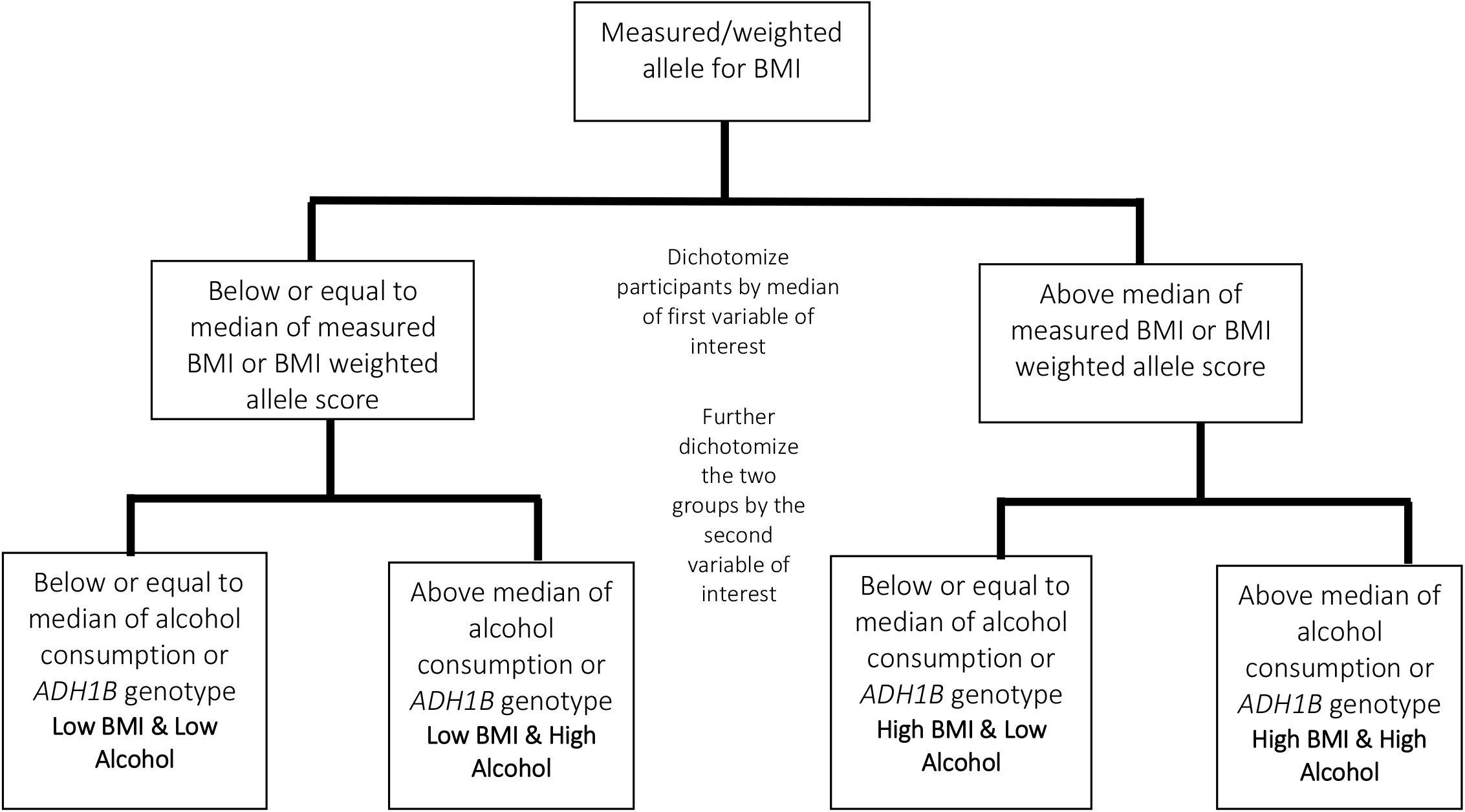
Flow diagram for the generation of factorial groups, using observational and genetic data. **Factorial multivariable**: Both body mass index (BMI) and weekly alcohol consumption were dichotomized based on the median of measured or self-reported values. Those equal to, or below the median were categorised as in the low group, whilst those above the median were categorised in the high group. **Factorial Mendelian randomisation (MR)**: For genetic propensity, BMI was categorised according to the median of the weighted allele score for BMI, where those equal to or below the median were categorised as having low BMI, and those above the median were categorised as having high BMI. Alcohol propensity was determined according to ADH1B alleles. Individuals that were homozygous for the alcohol decreasing traits and heterozygous individuals were combined, as determined to be appropriate from previous MR analyses of these traits on alcohol intake (11), to create the low alcohol propensity group. The high alcohol propensity group contains all individuals homozygous for the alcohol increasing trait. Figure adapted from (21)

The aim of this study was to use observational regression and factorial MR analyses to investigate the joint effect of BMI and alcohol consumption on liver injury biomarkers (ALT and GGT) and on liver disease.

## Methods

We used data from the Copenhagen General Population Study (N=98 643), a large population cohort which has collected genotypic and phenotypic data of relevance to a wide range of health-related problems. Residents of Copenhagen, aged 20 or older and of white, Danish descent were randomly selected from the national Danish Civil Registration for recruitment between 2003 and 2014. Ethics approval for the study was obtained from Herlev Hospital and Danish ethical committee and all participants provided written informed consent. Additional study details have been previously published (9).

Genetic data were available on 96 185 (98%) participants. BMI and alcohol data were missing for a further 387 individuals, leaving 95 798 (97%) participants available for analysis. There was some further missing data for other covariables. A diagnosis of liver disease is likely to result in treatment and lifestyle changes, therefore individuals with known liver disease at baseline (prevalent cases) were excluded from our main analyses (**Figure 2**).

**Figure 2:**
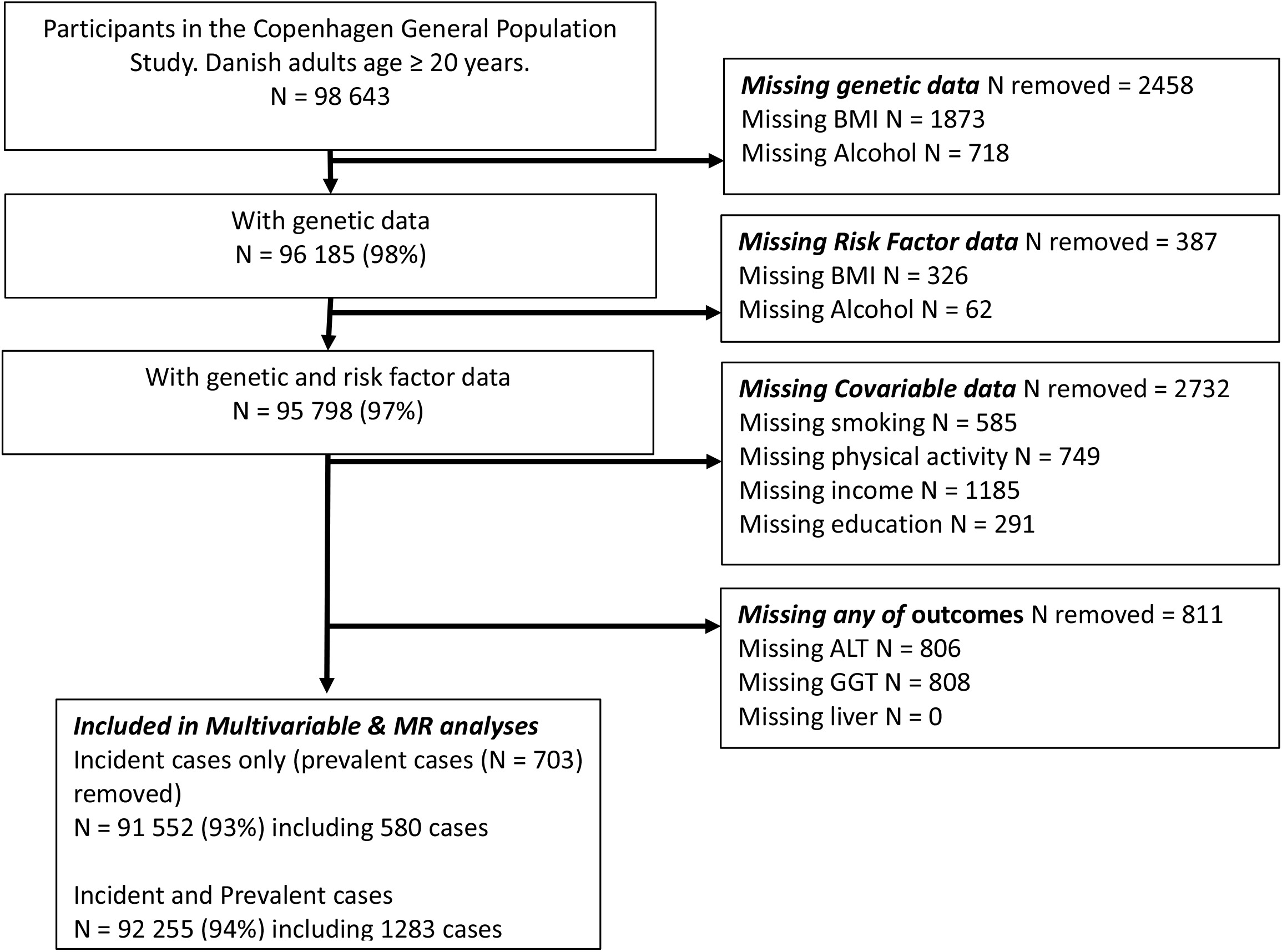
Missing data and total number of participants for each analysis.

All measurements were completed by trained staff at one clinic centre. Weight was measured without shoes and in light clothing to the nearest 0.1 kg on Soehnle Professional scales. Height was measured to the nearest 0.1 cm with a Seca stadiometer. Usual alcohol intake was reported as weekly consumption of beer in bottles and standard glasses of wine and spirits. Each of these in Denmark contains the equivalent of ~12g of pure alcohol, equal to 1.5 units of alcohol (a unit being 8g of alcohol).

We used five genetic variants that are robustly associated with BMI and fat mass to create a weighted allele score as an instrumental variable for BMI: *FTO* rs9939609, *TMEM18* rs6548238, *MC4R* rs17782313, *BDNF* rs10767664 and *GNPDA2* rs6548238. This score was externally weighted according to the beta for each genetic variant association with BMI from the most recent genome-wide association study (10). A single genetic variant, *ADH1B* (rs1229984, Arg47His in exon 3), was used as an instrumental variable for alcohol consumption. This is robustly associated with liver disease enzymes, as demonstrated in previous analyses using these data (9, 11). Genotyping was conducted by laboratory technicians without access to participant data, using the ABI PRISM 7900HT Sequence Detection System (Applied Biosystems Inc., Foster City, CA, USA), and TaqMan assays. Genotyping was verified by DNA sequence in at least 30 individuals with each genotype. Reruns were performed twice and 99.96% of all available participants were genotyped.

Plasma ALT and GGT were measured using standard hospital assays (Konelab and ACL) and were subject to daily internal quality control assessing assay precision and monthly external quality control assessing assay accuracy. Participants were linked to their hospital records providing information on all admissions and outpatient appointments prior to and after study recruitment. These identified individuals with existing liver disease and those who developed liver disease during follow-up (incident cases). Participants were also linked to death records. The following ICD diagnostic terms were included as cases of liver disease: cirrhosis including alcoholic fatty liver, unspecified chronic liver disease without mention of alcohol, hepatic fibrosis, malignant neoplasms of the liver, acute and subacute necrosis of the liver, chronic passive congestion of the liver – unspecified disorder of the liver, abdominal swelling, other non-specific liver disease. **sTable 1** provides the ICD codes used.

Participants completed a questionnaire that recorded details on smoking, physical activity, income and educational attainment, which could be included as covariates.

### Statistical analyses

All analyses were conducted in Stata version 14.

We explored the separate and joint effects of BMI and alcohol on ALT, GGT and liver disease using both observational multivariable and MR analyses. In multivariable and MR analyses, the results reflect the same magnitude of change in exposures (i.e. 1 SD-score greater BMI or 1-unit greater alcohol intake on outcome). ALT and GGT were natural log-transformed to normalise the distribution of residuals in regression models. BMI was standardised for age (5-year groups) and sex into standard deviation (SD) scores. Both the MV and MR analyses are on a multiplicative scale and results for biomarkers are presented as percentage differences in mean levels per 1 BMI SD or 1 unit of alcohol.

### Multivariable analyses

In multivariable analyses we examined BMI in categories of normal/underweight (<25 kg/m^2^), overweight (25-29.9kg/m^2^) and obese (≥30kg/m^2^), and as a continuous BMI SD-score. We examined alcohol in categories of none, 1-14, and ≥ 15 units per week, and per unit of alcohol consumption per week. We used multivariable linear regression to examine associations with ALT and GGT and logistic regression for associations with liver disease.

To examine joint associations of BMI and alcohol with markers of liver injury, a factorial analysis was completed, by categorising participants in four groups according to their measured values of the exposure: high BMI/high alcohol (reference), high BMI/low alcohol, high alcohol/low BMI and low BMI/low alcohol. Low and high BMI were defined as ≤ or > median BMI SD-score, respectively. Similarly, low and high alcohol were defined as ≤ or > median alcohol consumption, respectively. The mean difference between high vs low groups was 1.49 units for BMI and 14.68 units/week for alcohol (**Figure 1**). Multivariable linear or logistic regression was then used to assess the association of each of these groups with outcomes. Stratified analyses of BMI SD-score with each outcome by strata of alcohol consumption (none, < 15 and ≥ 15 units per week) were also performed. A likelihood ratio test was used to test for statistical evidence of interaction by comparing a model with BMI SD-score and alcohol categories as independently to one where an interaction term was included.

For all multivariable analyses we considered the ‘best’ estimate of a causal effect to be adjusted for all observed variables that we considered potential confounders (age, sex, cigarette smoking, physical activity, educational attainment and income) and individuals with baseline liver disease removed.

### Mendelian randomization analyses

An exact test was used to examine Hardy-Weinberg equilibrium of genotype frequencies (12). We present the first-stage (regression of risk factor on genetic instrument) F-statistic and R^2^ as indicator of the strength of the association between genetic instruments and observed phenotype. We measured associations of BMI and alcohol instruments with observed confounders to test our *a priori* assumption of no confounding associations. MR analyses were completed using the 2-stage least squares instrumental variable regression method for continuous outcomes. For the binary outcome (liver disease), MR was run in two regression stages, including robust standard errors in the second stage (13, 14).

As with multivariable analyses a factorial analysis was completed to test joint associations, categorising participants to the same four groups previously described according to genetic propensities. Low and high BMI were defined as ≤ or > median BMI weighted allele score, respectively. Alcohol was dichotomized according to *ADH1B* alleles, where individuals homozygous for the alcohol decreasing allele or heterozygous for *ADH1B* were classed as low alcohol, whilst the high for alcohol group were individuals homozygous for the alcohol increasing allele of *ADH1B* (11) (**Figure 1**). The mean difference between high vs low groups was 0.5 BMI SD and 1.8 units/week for alcohol (**Figure 1**). Regression methods were then used to assess the association of each of these groups with outcomes. We investigated evidence of an interaction between BMI and alcohol using stratified MR of the BMI weighted allele score instrument by self-reported weekly alcohol consumption (none, < 15 and ≥ 15 units/week). A variance weighted least squares approach was used to test for statistical evidence of a difference in circulating biomarkers and liver disease risk across alcohol strata. This was done by calculating a chi squared test to assess for deviations between point estimates in each stratum.

### Comparing multivariable and MR results

We compared the multivariable and MR results by using a chi-squared test to test the null hypothesis that the coefficients are consistent with one another. We made this comparison for analyses of the association of BMI and alcohol separately, and for the stratified results of the effect of 1SD greater BMI within strata of reported alcohol consumption. We were not able to compare the magnitude of effect sizes of the multivariable and MR factorial analyses because it was not possible to split participants into high and low BMI and alcohol groups in identical ways for the two methods. Thus, in the multivariable analyses, the high versus low results reflect a 1.5 SD difference in BMI and a 14.7 difference in units of weekly alcohol consumption, whereas the equivalent differences in the MR analyses are 0.5 SD and 1.8 units per week, respectively. We do, however, compare the directions and overall patterns of association/effects for the factorial results.

### Sensitivity analyses

Sensitivity analyses were completed to test the robustness of our MR results for the effect of BMI on outcomes. These included using methods that are more robust to pleiotropic variants (i.e. weighted median methods and MR-Egger method) (15, 16), and assessing the impact of outlying variants by removing one variant at a time and recalculating the overall MR estimate (i.e. leave-one-out analysis). As we have only one genetic variant for alcohol these sensitivity analyses were not possible. For MR analyses of both BMI and alcohol we adjusted for income and smoking because of some evidence of genetic instrument associations with these as a sensitivity analysis. All analyses were repeated with individuals with prevalent cases of liver disease included. Full details of these analyses are available in the **Supplementary material**.

## Results

Of the 98 643 participants, 9% reported not drinking alcohol and 12% reported drinking 22 or more units per week (**sTable 2**). Forty-three percent of participants were normal-weight, 1% underweight, 40% overweight and 16% obese. There were 1405 (1.4%) cases of either prevalent or incident liver disease, with 616 (0.6%) incident cases. Alcohol intake and BMI differed by gender (**sTable 2**). All measured confounders were associated with reported alcohol intake and measured BMI (sTable 3A) but most were not associated with *ADH1B* or the BMI weighted allele score (**sTable 3B**). There was some evidence that BMI weighted allele score was positively associated with smoking and negatively associated with income. All observed confounders were associated with the factorial groups in multivariable analysis; whilst none were associated with factorial groups in MR analysis (**sTable 3C**). The minor allele frequencies for all five genotypes are shown in **sTable 4**; all were in Hardy-Weinberg equilibrium. The first stage partial F-statistic and R^2^ for the combined weighted allele score for BMI were 453 and 0.0049, and for *ADH1B* were 117 and 0.0013, respectively (**sTable 5**).

In both multivariable and MR analyses, individually higher BMI and higher alcohol consumption were associated with higher mean ALT, GGT and odds of incident liver disease (**Figure 3**). Overall, multivariable and MR results were consistent with each other, although point estimates for ALT and GGT appeared slightly larger in multivariable compared with MR analyses (**Figure 3A and 3B**). For both methods the magnitudes of the associations with ALT and GGT were similar for BMI, whereas alcohol was more strongly (more than double the magnitude) associated with GGT than ALT.

**Figure 3:**
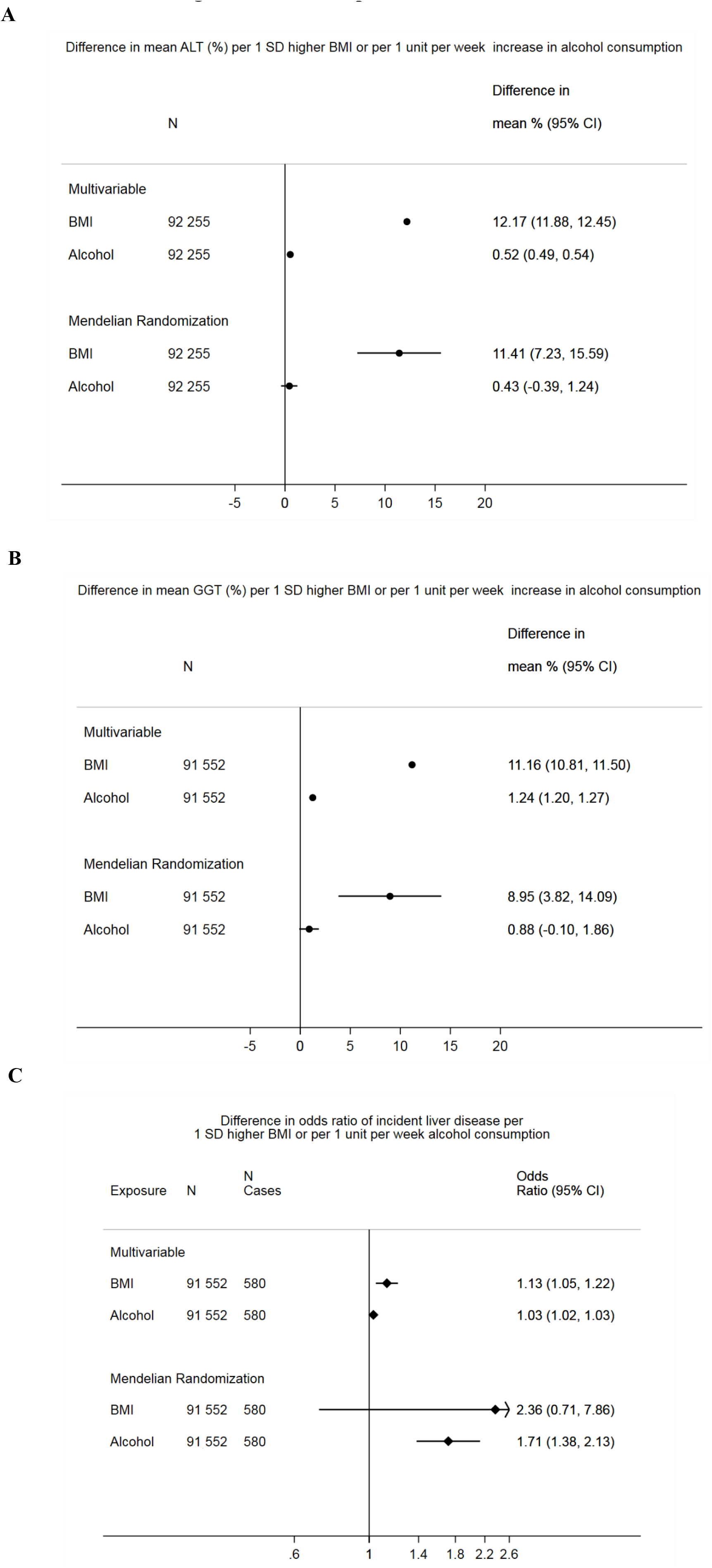
Multivariable adjusted and Mendelian randomization analyses of the individual effects of body mass index (BMI) and alcohol on liver injury enzymes (A) alanine aminotransferase (ALT), (B) γ-glutamyltransferase (GGT) and (C) incident liver disease, excluding individuals with prevalent liver disease. BMI measured as age and sex standard deviation (SD) units. Alcohol measured as units of alcohol consumed per week, where one unit is equivalent to 12g of alcohol (SD = 10.22). Multivariable analyses are adjusted for age, sex, smoking, education, income and physical activity. Mendelian Randomization using weighted allele score for BMI and *ADH1B* alleles for alcohol consumption. CI = confidence interval

Stronger effects of both BMI and alcohol were seen in the MR analysis compared with multivariable regression for incident liver disease. However, the confidence intervals in the MR analyses were very wide and included the null value, particularly in the case of BMI (**Figure 3C**). Analysis including individuals with prevalent cases of liver disease showed similar effect estimates to analyses restricted to incident cases (**sFigure1 and sFigure2**)

Multivariable analysis indicated a linear relationship of increasing BMI and increasing alcohol consumption with increasing levels of biomarkers for liver injury and odds of disease (**sFigure3 & sFigure4;** non-linear associations were not tested in MR analyses).

In both multivariable and MR factorial analyses, those in the low BMI and low alcohol consumption groups had the lowest mean ALT and GGT in comparison to those in the highest categories of both, with the low BMI/high alcohol and high BMI/low alcohol groups being in between these two (**Figure 4A-D**). In both sets of analyses, being in the low BMI and high alcohol group conferred a greater protective effect on ALT than those in the high BMI and low alcohol group compared to the reference group. However, for GGT being low for either BMI or alcohol had almost identical protective effects compared with the reference group, in both multivariable and MR analyses. These results for liver biomarkers were very similar when individuals with prevalent cases of liver disease were included in the analysis (**sFigure5**).

**Figure 4:**
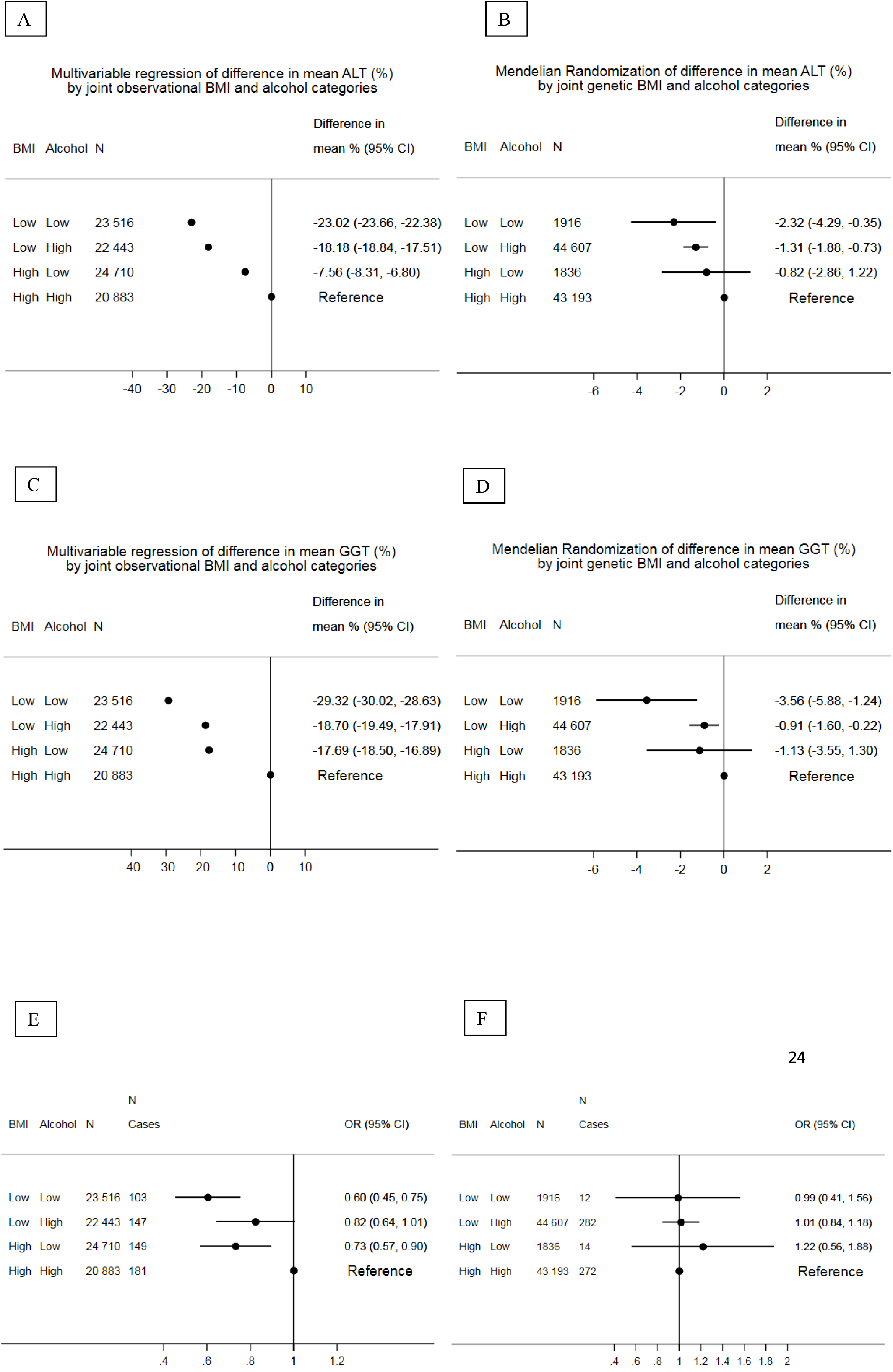
Multivariable and Mendelian randomization factorial analyses assessing the joint effects of body mass index (BMI) and alcohol on biomarkers of liver injury and incident liver disease. Low vs high BMI = 1.49 km/m^2^ difference in MV analyses and 0.51 km/m^2^ in MR analyses. Low vs high alcohol = 14.68 units per week difference in MV analyses and 1.78 units per week difference in MR analyses, where one unit of alcohol is equivalent to 12g. MV = Multivariable analysis, adjusted for age, sex, smoking, education, income and physical activity. MR= Mendelian Randomization CI = confidence interval. ALT = Alanine Amino Transferase. GGT = γ-glutamyltransferase. BMI = Body mass index

For odds of incident liver disease, factorial multivariable regression analyses had a similar pattern to the biomarkers (i.e. reduced odds in those low for both BMI and alcohol, in comparison to those high for both, and in between for the two mixed groups) (**Figure 4E-F**). This pattern was not consistent in the factorial MR analyses, where the odds ratio for being low for both BMI and alcohol or low for BMI and high for alcohol was one. The group with high BMI and low alcohol consumption appeared to have greater odds of liver disease compared to those in the group with high BMI and high alcohol intake. Although in all groups the confidence intervals in the MR factorial analyses were wide and included the null. When analyses were repeated with both incident and prevalent liver disease included as the outcome, factorial multivariable analyses suggested that those with low BMI and high alcohol had similar reduced odds to those with low values of both risk factors, which was not confirmed by factorial MR (**sFigure6**).

In stratified multivariable analyses, the positive association of BMI with all outcomes was strongest in those consuming 15 or more units of alcohol per week compared with those not drinking alcohol or drinking less than 15 units per week. There was strong statistical support for a positive interaction on a multiplicative scale between the two risk factors for ALT and GGT; this evidence was weaker for odds of liver disease, but the pattern of interaction was similar (**sFigure 7**). The same pattern of positive multiplicative interaction of BMI and alcohol was not observed for any of the outcomes in MR analyses (**sFigure 7**). For ALT and GGT the MR results suggested a weaker positive effect of BMI in those with highest alcohol consumption (15 units or more per week), although these were imprecisely estimated, and we found no strong evidence for any interaction. For liver disease, the stratified MR analyses were particularly imprecise, though again results suggested, if anything a negative interaction.

### Sensitivity analyses

Weighted median MR and MR Egger estimates for the effect of BMI on outcomes were broadly similar to our main MR analyses results (**sTable 6**). Heterogeneity between the five BMI related genetic variants used as instrumental variables in MR was modest (I^2^ = 52%, 13% and 38% for the effect of BMI on ALT, GGT and odds of incident liver disease). Results were consistent when one genetic variant at a time was removed, although in both ALT and GGT the estimates were attenuated slightly when *FTO* was removed (**sFigure 8**). Adjustment of the MR analyses for smoking and income did not notably influence the results.

## Discussion

Using both multivariable and MR analyses we have shown that higher BMI and alcohol consumption are important causal risk factors for higher circulating GGT and ALT and odds of incident liver disease. The broad consistency of findings between multivariable and MR analyses for each exposure on ALT and GGT provides more compelling evidence for causality than either method alone. They each have different key sources of bias, and it would be unlikely to observe similar results if these were due to residual confounding in the multivariable analyses and pleiotropy in the MR analyses (17). MR findings for incident liver disease suggested a stronger effect of both BMI and alcohol than the observational multivariable findings, suggesting the possibility of masking residual confounding for this outcome in multivariable analyses or horizontal pleiotropy exaggerating effects in MR analyses. However, MR estimates for incident liver disease were imprecise and the wide-confidence intervals for BMI included the null value.

Consistent with individual effects, multivariable and MR factorial analyses suggested that adults with low BMI and low alcohol consumption had lower ALT and GGT than those with high levels of both risk factors. Those with low levels of one risk factor had intermediate levels. The higher concentration of ALT and GGT in the high BMI/high alcohol group could reflect the combination of independent effects of BMI and alcohol or a positive interaction between the two risk factors. We found some evidence of positive multiplicative interactions of BMI and alcohol with ALT and GGT in multivariable analyses but not in MR analyses. For incident liver disease, the multivariable factorial and stratified analyses followed the same pattern of those seen with ALT and GGT, whereas in MR analyses there was no clear evidence of differences across factorial groups. However, confidence intervals for factorial or interaction analyses for incident liver disease were wide, suggesting lack of statistical power despite the large sample size.

The differences in the stratified analyses between multivariable and MR, could reflect multivariable results being biased by confounders that act differently within strata of alcohol consumption or residual confounding not measured here. Lack of statistical power in the MR analyses may also be important. Given the consistency of findings between the multivariable and MR results for the individual and joint effects of BMI and alcohol on ALT and GGT, we consider our results to provide most support for BMI and alcohol acting independently of each. However, we do not see consistent results between multivariable and MR analyses for liver disease as the outcome, where the MR factorial analysis suggests there is little difference in risk of liver disease between the BMI and alcohol groups. Although this is possibly due to small numbers of cases, it could be a true effect of an inhibitory interaction.

Our multivariable analyses, are consistent with most previously published studies of the individual effects of BMI and alcohol on liver biomarkers and disease. Using MR, in a smaller number of participants from the same study population as used here (N = 58 313) we have shown a causal effect of greater alcohol consumption on ALT and GGT (11). We have added to that previous work by estimating the causal effect of increasing BMI on ALT, GGT, and incident liver disease and undertaking the first, to our knowledge, MR analyses of the joint effects of BMI and alcohol on these outcomes. Consistent with our multivariable analyses, some (5, 6), though not all (7), previous multivariable studies report a positive multiplicative interaction between higher BMI and alcohol consumption on liver biomarkers and disease. However, our MR analyses suggest that BMI and alcohol combine as we would expect if their individual effects are independent of each other on ALT and GGT, as we observe a non-linear pattern in stratified MR, and possibly with weaker effects than independence for liver disease, though power will have been low. In the absence of other MR studies of these joint effects we would suggest caution regarding the joint effects with liver disease outcomes.

### Strengths and limitations

Our study has several strengths, including the large sample size, the assessment of potential single and joint causal effects of BMI and alcohol on liver outcomes using both multivariable and MR, appropriate sensitivity analyses to test the robustness of the MR assumptions and assessing effects on liver disease outcomes (as well as biomarkers), which, because they are relatively rare, has not been explored previously with MR, or in many multivariable regression studies. The study of liver biomarkers, high levels of which reflect cell damage in the liver and biliary track, as well as disease outcomes, is valuable for indicating propensity to future disease and allowing analyses with greater statistical power.

MR and multivariable analyses are potentially biased by violation of their assumptions. Multivariable approaches assume that all confounders have been accurately measured and accounted for, there is no reverse causation, selection bias or systematic measurement error that distorts the estimate (17). The removal of those with prevalent liver disease in our main analyses should limit potential survivor bias and reverse causality. We adjusted for key observed potential confounders, but residual confounding by unmeasured confounders or imprecise measurement, might bias causal estimates in our multivariable analyses. For the individual effects of BMI and alcohol on ALT, GGT and liver disease multivariable results were similar to the MR results, which are less prone to confounding, suggesting that residual confounding might not be a major issue here (18). Though the stronger individual effects in MR analyses of liver disease might mean that multivariable results were biased by masking residual confounding.

MR analyses assume that there is a robust association between the genetic instrument and risk factor, that the genetic instrument is not associated with confounders of the risk factor-outcome association and that there is no influence of the genetic instrument on the outcome other than via the risk factor of interest (the latter may be violated by horizontal pleiotropy) (19, 20). We selected genetic variants for both BMI and alcohol that have been shown to be robustly (including replicating across many studies) related to these outcomes, and first-stage F-statistics and R^2^ for both instruments were high. We did find some association of BMI genetic variants with smoking and income, but adjustment for these in the MR analyses did not alter results. The Copenhagen General Population Study participants are ethnically and geographically homogenous, which reduces the risk of confounding due to population stratification. Our sensitivity analyses, suggest some greater influence of one of the BMI genetic variants *FTO* than other variants but the results are not markedly different, and our conclusions are not altered, by its removal.

An important limitation of both analyses is that those reporting no consumption of alcohol may be a heterogeneous group of life-long abstainers and those who have previously consumed alcohol but have since become abstainers. However, the multivariable regression results suggest linear associations of increasing consumption from this group of non-drinkers and the similarity of analyses including and excluding prevalent disease cases suggest that this is unlikely to have introduced major bias. Despite our large sample size our MR analyses are less precisely estimated than the multivariable results.

## Conclusion

Taking account of both our multivariable and MR results this study suggests that high BMI and alcohol consumption act independently to increase biomarkers of liver injury and risk of liver disease. Interventions to reduce both of these risk factors might produce the greater benefit in terms of reducing population levels of biomarkers of liver injury than interventions aimed at either BMI or alcohol alone. However, the current evidence is not clear whether this will directly translate to a reduced risk in clinical liver disease and further studies are required to further identify the true joint causal effect of BMI and alcohol on liver disease.

## Acknowledgements

We thank all of the study participants.

## Authors’ contributions

Study concept and design: Lawlor, Borges, Nordestgaard, Davey Smith;

Acquisition of data: Nordestgaard, Tybjærg-Hansen, Benn;

Database handling and updating: Benn;

Statistical analysis plan: Carter, Borges, Lawlor;

Completing of statistical analyses: Carter;

Drafting of the manuscript: Carter, Borges, Lawlor;

Interpretation of results: Carter, Borges, Lawlor;

Critical revision of the manuscript for important intellectual content: Carter, Borges, Lawlor, Nordestgaard, Benn, Tybjærg-Hansen, Davey Smith;

Obtained funding: Nordestgaard, Tybjærg-Hansen;

Administrative, technical, or material support: Nordestgaard, Benn;

Carter, Borges, Lawlor, Nordestgaard and Benn had full access to all of the data in the study and take responsibility for the integrity of the data (Nordestgaard and Benn) and the accuracy of the data analysis (Lawlor, Carter and Borges).

## References

1. Bellentani S, Marino M. Epidemiology and natural history of non-alcoholic fatty liver disease (NAFLD). Ann Hepatol. 2009;8 Suppl 1:S4–8.

2. Corey KE, Kaplan LM. Obesity and liver disease: the epidemic of the twenty-first century. Clin Liver Dis. 2014;18(1):1–18.

3. Bosetti C, Levi F, Lucchini F, Zatonski WA, Negri E, La Vecchia C. Worldwide mortality from cirrhosis: an update to 2002. J Hepatol. 2007;46(5):827–39.

4. Hart CL, Morrison DS, Batty GD, Mitchell RJ, Davey Smith G. Effect of body mass index and alcohol consumption on liver disease: analysis of data from two prospective cohort studies. BMJ. 2010;340:c1240.

5. Alatalo PI, Koivisto HM, Hietala JP, Puukka KS, Bloigu R, Niemela OJ. Effect of moderate alcohol consumption on liver enzymes increases with increasing body mass index. Am J Clin Nutr. 2008;88(4):1097–103.

6. Puukka K, Hietala J, Koivisto H, Anttila P, Bloigu R, Niemela O. Additive effects of moderate drinking and obesity on serum gamma-glutamyl transferase activity. American Journal of Clinical Nutrition. 2006;83(6):1351–4.

7. Liu B, Balkwill A, Reeves G, Beral V, Million Women Study C. Body mass index and risk of liver cirrhosis in middle aged UK women: prospective study. BMJ. 2010;340:c912.

8. Ference BA, Majeed F, Penumetcha R, Flack JM, Brook RD. Effect of naturally random allocation to lower low-density lipoprotein cholesterol on the risk of coronary heart disease mediated by polymorphisms in NPC1L1, HMGCR, or both: a 2 × 2 factorial Mendelian randomization study. J Am Coll Cardiol. 2015;65(15):1552–61.

9. Lawlor DA, Nordestgaard BG, Benn M, Zuccolo L, Tybjaerg-Hansen A, Davey Smith G. Exploring causal associations between alcohol and coronary heart disease risk factors: findings from a Mendelian randomization study in the Copenhagen General Population Study. Eur Heart J. 2013;34(32):2519–28.

10. Locke AE, Kahali B, Berndt SI, Justice AE, Pers TH, Day FR, et al. Genetic studies of body mass index yield new insights for obesity biology. Nature. 2015;518(7538):197–206.

11. Lawlor DA, Benn M, Zuccolo L, De Silva NM, Tybjaerg-Hansen A, Smith GD, et al. ADH1B and ADH1C genotype, alcohol consumption and biomarkers of liver function: findings from a Mendelian randomization study in 58,313 European origin Danes. PLoS One. 2014;9(12):e114294.

12. Cleves M. Hardy-Weinberg Equilibrium Tests and Allele Frequency Estimation. STATA Technical Bulletin. 1999:34–7.

13. Christopher F Baum MES, Steven Stillman. IVREG2: Stata module for extended instrumental variables/2SLS and GMM estimation. Statistical Software Components. 2002.

14. Palmer TM, Holmes MV, Keating BJ, Sheehan NA. Correcting the Standard Errors of 2-Stage Residual Inclusion Estimators for Mendelian Randomization Studies. Am J Epidemiol. 2017;186(9):1104–14.

15. Bowden J, Davey Smith G, Burgess S. Mendelian randomization with invalid instruments: effect estimation and bias detection through Egger regression. Int J Epidemiol. 2015;44(2):512–25.

16. Bowden J, Davey Smith G, Haycock PC, Burgess S. Consistent Estimation in Mendelian Randomization with Some Invalid Instruments Using a Weighted Median Estimator. Genet Epidemiol. 2016;40(4):304–14.

17. Lawlor DA, Tilling K, Davey Smith G. Triangulation in aetiological epidemiology. Int J Epidemiol. 2016;45(6):1866–86.

18. Smith GD, Lawlor DA, Harbord R, Timpson N, Day I, Ebrahim S. Clustered environments and randomized genes: a fundamental distinction between conventional and genetic epidemiology. PLoS Med. 2007;4(12):e352.

19. Lawlor DA, Harbord RM, Sterne JA, Timpson N, Davey Smith G. Mendelian randomization: using genes as instruments for making causal inferences in epidemiology. Stat Med. 2008;27(8):1133–63.

20. Davey Smith G, Hemani G. Mendelian randomization: genetic anchors for causal inference in epidemiological studies. Hum Mol Genet. 2014;23(R1):R89–98.

21. Ference BA, Robinson JG, Brook RD, Catapano AL, Chapman MJ, Neff DR, et al. Variation in PCSK9 and HMGCR and Risk of Cardiovascular Disease and Diabetes. N Engl J Med. 2016;375(22):2144–53.

